# Sex Differences in Predictors of Seizure in Contrast-Enhancing Gliomas at Clinical Presentation: A Network Approach

**DOI:** 10.1101/708032

**Authors:** Sandra K Johnston, Aditya Khurana, Paula Whitmire, Sara Ranjbar, Akanksha Sharma, Andrea Hawkins-Daarud, Joshua B Rubin, Alyx B Porter, Peter D Canoll, Kathleen Egan, Leland S Hu, Maciej M Mrugala, Priya Kumthekar, Kristin R Swanson

## Abstract

**Background:** Brain tumor related epilepsy (BTE) is a major co-morbidity related to the management of patients with brain cancer. Despite published practice guidelines recommending against anti-epileptic drug (AED) utilization in patients with gliomas, there is heterogeneity in prescription practices of AEDs in these patients. In an attempt to impact BTE management, we statistically analyzed clinically relevant attributes (sex, age, tumor size, tumor growth kinetics, and tumor location) pertaining to seizure at presentation and used them to build a computational machine learning model to predict the probability of a seizure (at presentation).

**Methods:** From our clinical data repository, we identified 223 patients (females, n=86; males, n=137) with pathologically-determined glioma and known seizure status at clinical presentation. Non-parametric and Fisher’s Exact tests were used to identify statistical differences in clinical characteristics. We utilized a random forest machine learning method for generating our predictive models by entire cohort and separated by male and female.

**Findings:** Patients were divided into those that presented with seizure (SP, n=96, 43%; F, n= 28; M, n= 68) and those that presented without seizure (nSP, n=127, 57%, F n=58, M n=69). Females presented with seizures significantly less often than males (x^2^=6·28, p=0·01). SP patients had significantly smaller T1Gd radius compared to nSP (SP 11·30mm, nSP 18.66mm, p<0·0001). Tumor size and patient age were significant negative predictors for SP; patients with larger tumors, older age and less tumor diffusivity (p/D) were at lower risk for SP.

**Interpretation:** Despite heterogeneity across our patient cohort, there is strong evidence of a role for patient sex, tumor size, tumor invasion, and patient age in predicting the incidence of seizures at diagnosis. Future studies, with prospectively detailed data collection, may provide clearer insights into the incidence of seizures through a patient’s treatment course.

## Introduction

Brain tumor related epilepsy (BTE) presents a very different degree of complexity than that of idiopathic epilepsy, thus demanding its own body of research and exploration. Approximately one-third of patients with primary brain tumors will present with seizures and the lifetime risk of BTE ranges between 20 to 80%^1,2^. Rates of seizures are higher in lower grade gliomas (LGG, 40-80%) than higher grade gliomas (HGG, 30-50%)^3^. Tumors in the frontal, temporal, insular and centro-parietal regions are commonly associated with symptomatic seizures^4^. Yet, frequently, it is found that tumor location does not appear to be the same as the seizure focus^5^. Epidemiological studies have generally indicated overall incidence of epilepsy to be slightly higher in males than females, but a higher incidence of idiopathic generalized epilepsy has been seen in females in some studies^6^.

Understanding the epileptogenic nature of brain tumors will help us better identify who will have a seizure, and in turn allow us to optimize care. The involvement of cortical areas by tumor infiltration and growth likely contributes to the epileptogenic nature of certain tumors^7^. An increase in alkaline pH in the peritumoral space can induce excitatory pathways^8^. Elevation in extracellular glutamate contributes to epileptogenesis^9^, and glutamine synthetase levels have been linked to increased seizures in the glioma population. More recently, mutations in isocitrate dehydrogenase (IDH) have thought to result in the production of 2-hydroxyglutarate (2-HG) which in turn again increases the excitability of the environment (similar to glutamate)^10^. LGG are more likely to demonstrate an IDH mutation, and this association suggests a possible explanation for the higher rate of seizures in this type of brain tumor.

Despite the growing body of research on the etiology of BTE it remains less clear which gliomas will beget seizure at time of clinical presentation. This uncertainty often results in prophylactic administration of AEDs even though the American Academy of Neurology (AAN) issued a practice parameter in 2000 that recommended against this practice^1^. Despite the issuance of this guidance, actual practice varies significantly^11^. Furthermore, a 2004 meta-analysis showed that the resulting side effects of AEDs, including serious rash, sedation and coordination issues of several of the commonly used AEDs, outweighed the benefits of prophylactic use^12^. Drug interactions with chemotherapy agents are another significant concern with AED use^1^, side effects of AEDs often intensify when combined with chemotherapy and/or radiation therapy^13^. Without evidence demonstrating true efficacy, prophylactic AED usage cannot be well rationalized given the concern for side effects or poor health outcomes^13^.

Clinicians, especially those in neuro-oncologic and neuro-surgical practices, regularly face the challenge of AED prescription determination in patients with newly identified brain mass and would greatly benefit from having concrete predictors of which patients are more likely to experience BTE. The aim of this project was to assess which clinical variables were predictive of seizure at presentation in order to provide guidance for clinicians on the use of AED.

## Methods

### Patient Cohort

Our Institutional review board (IRB) approved data repository of clinical and imaging data is comprised of patients diagnosed with glioma that were either consented prospectively or approved for retrospective research. We selected patients from the repository based on the following eligibility criteria: a) pathological diagnosis of glioma (all grades, I-IV), b) contrast-enhancing tumor (measureable T1Gd abnormality) and c) known seizure-presentation status at time of initial tumor presentation. Clinical data, including sex, was acquired from medical record reports. Overall survival (OS) was calculated from the time of initial surgical intervention to the time of death. Patients who are still alive or are lost-to-follow-up were censored on their last date known alive. Histological grading of tumors was acquired either from pathology reports or progress notes in the absence of pathology reports.

Tumor location was acquired through image and medical record review. Lobe location was described as temporal, parietal, occipital, and/or frontal. Tumors located in more than one lobe were categorized as “multi-lobular,” further we counted the number of lobes containing tumor. Laterality of tumors was categorized into left-side, right-side or bilateral (crossing the corpus callosum and/or presence of tumors on both sides of the corpus callosum).

Seizure presentation status at time of tumor presentation on MR was abstracted by medical record review for mention of seizure activity. While EEG is a frequent part of routine epilepsy workup, in BTE it is less frequently obtained given it is not needed for the diagnosis, which can be made on clinical history alone. In our cohort pre-operative EEG was acquired only for seven patients (four Seizure Presenting (SP), three non-Seizure presenting (nSP)).

### Tumor Size and Image-Derived Growth Rates

Tumor size was determined from pre-treatment MR images including T2-weighted (T2) or T2 fluid-attenuated inversion recovery (T2-Flair) sequence and T1-weighted post-gadolinium contrast agent (T1Gd) sequence. The tumor-associated abnormality was segmented by trained individuals using our in-house threshold-based semi-automatic segmentation software on both T1Gd and T2 or T2-FLAIR (T2/FLAIR). These segmentations are used to calculate a tumor volume, which is converted to the spherically-equivalent radius for analysis (T1Gd radius and T2/FLAIR radius).

Using these values and our lab’s proliferation-invasion (PI) mathematical model of the kinetics of tumor growth we estimated a patient-specific invasiveness ratio (p/D) for each patient^14^. The PI mode is based on the two hallmark features of glioma growth: proliferation (p) and invasion (D); in this ratio, p(1/yr) is a net proliferation rate and D (mm^2^/yr) is a net migration rate. In order to obtain the full complement of model kinetics two pre-treatment MR image dates are necessary but are often not clinically acquired. Thus, we turned to our more advanced mathematical model, Proliferation-Invasion-Hypoxia-Necrosis-Angiogenesis (PIHNA), that extends the PI model by incorporating elements of angiogenesis, tumor hypoxia and necrosis^15^. PIHNA entails use of a look-up table with the relative size of necrosis used in lieu of velocity to estimate growth parameters and thus achieve measures of D and p using a single multi-modal MR image time point^16^.

## Statistical Analyses

We used statistical methods to broadly investigate the potential associations between the variables and seizure at presentation status. We used the Welch’s t-test (2-tailed, unpaired) for continuous numeric attributes, Fisher’s exact tests, and chi-squared tests. Values listed are mean unless otherwise specified; standard deviation (SD) is included as ± following each mean. Kaplan-Meier (K-M) test used for survival statistics, median values include interquartile range (IQR). Statistical significance was defined at p = 0.05 for all comparisons. As sex is an important determinant of outcome, we conducted sex differences analysis and reported where relevant^17^.

### Random Forest Predictive Modeling

#### Mixed Cohort Model

We explored the utility of random forest machine learning method in classifying SP and nSP cases. Using an 80-20 ratio, we used 176 patients (76 SP, 100 nSP) for training and the remaining 45 patients (19 SP, 26 NSP) were used for testing. The training set consisted of 65 females (37%) while the testing set consisted of 20 females (44%). To ensure optimal use of the sample size, we deferred from removing missing values and instead replaced them with median of the predictor across the training set. We ensured that the proportion of seizure-presenting and non-seizure presenting patients remained the same between training and test sets. We adjusted the hyper-parameters of RF using a grid search approach in which optimum combination of hyperparameters is found by assessing the performance of candidate models using all possible combinations of hyperparameters. The candidate with the highest performance on the validation set was selected as the optimal classifier. The performance of the selected model was measured using 10-fold cross validation scheme during training as well as validated on the separate test set. Performance measures were area under the ROC curve (AUC), accuracy, sensitivity, and specificity of predictions (average of folds in cross validation).

#### Sex Specific Models

Additionally, in order to remove potential sex biases in predictions, we split the cohort by patient sex and repeated train-test split. There were 136 males in the cohort, 108 (53 SP, 55 nSP) were used for training and 28 in testing (14 SP, 14 nSP). For the 85 females, we used 72 (21 SP, 49 nSP) for training and 15 (seven SP, eight nSP) for testing. We handled the class imbalance in female cohort by resampling the minority class and matching the size of majority class before training. Performance of the models was assessed similar to the mixed cohort model.

## Results

### Patient population

From our data repository, 223 patients (Table 1) met eligibility criteria (SP, 43% n=96; nSP, 57% n=127). Seizure semiology was available for 53 (55%) of SP patients (seizure type: generalized, n=25; partial complex, n=8; partial simplex, n=20). Presentation for those patients without seizure included headache, gait imbalance, memory decline, cognitive impairment, mental status changes, lack of coordination, speech impairment, visual changes, sensory involvement, and/or personality changes.

**Table 1.**
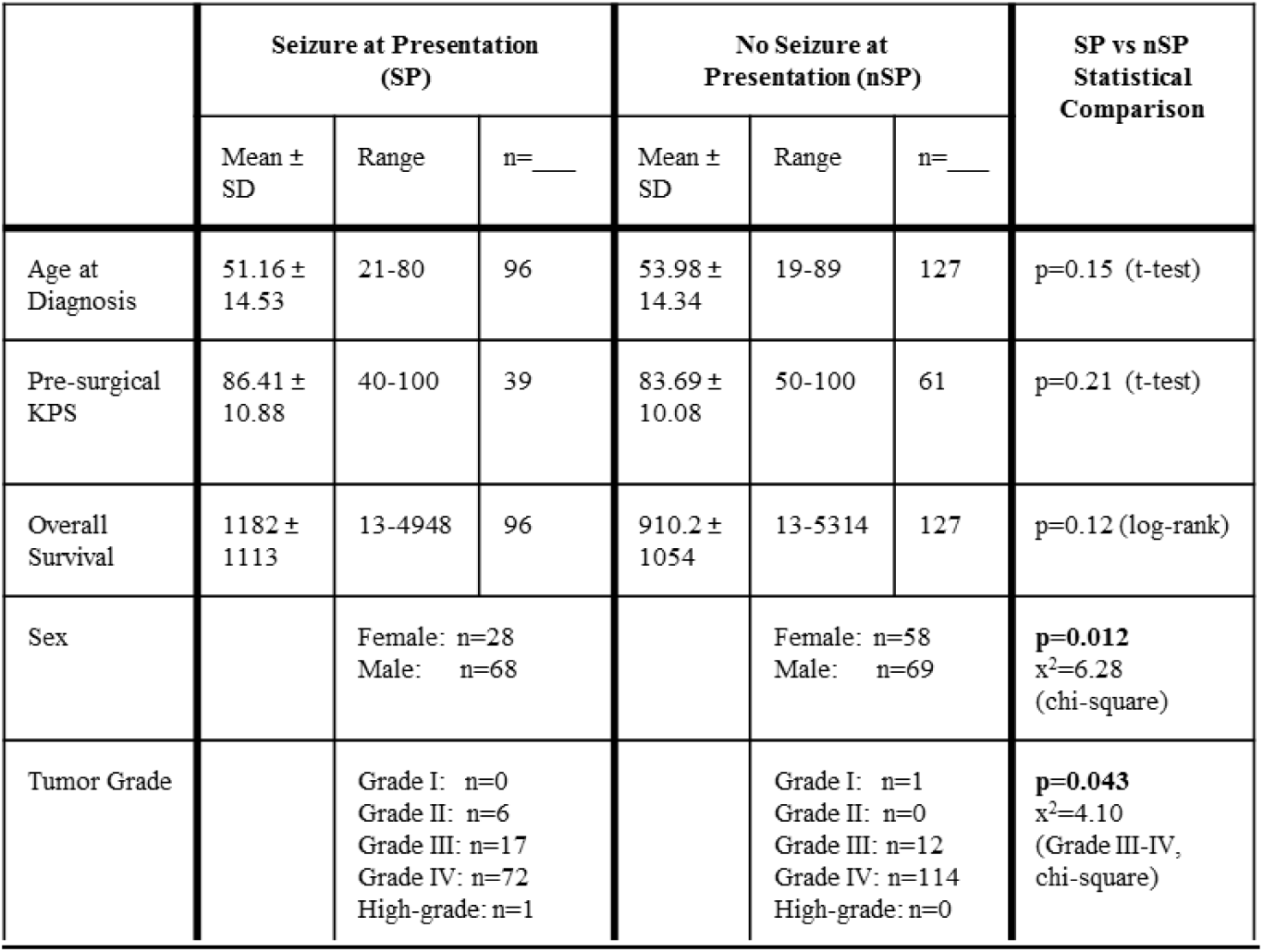
Subject Demographics and Tumor Characteristics. **Supplement Table S2** shows demographic breakdown by males and females.

Seizure presentation was not independent of patient sex, with females presenting with seizures significantly less often (32.6%: 28 SP, 58 nSP) than males (49.6%: 68 SP, 69 nSP) (x^2^=6·28, p=0.01). Overall, there was a M:F ratio of 2.4:1 among SP vs 1.9:1 among nSP. Furthermore, in males only there was a significant dependence between seizure presentation status and tumor grade (**Supplement Table S2**). The majority (97%) of our cohort were HGG, (Grades III and IV) presenting without seizure (59%). Amongst Grade IV gliomas females have a significantly (chi square 4.86, p=0.03) smaller proportion of SP to nSP (SP:nSP ratio: females 1:2.5 males 1:1.2 for males). Seizure presentation was more common among low grade tumors. All six grade II patients presented with seizures. When comparing grade III (17 SP vs 12 nSP) and grade IV (72 SP vs 114 nSP) patients, seizure presentation was found to be dependent on tumor grade (x^2^=4·10, p=0.04). There was no statistical significance in age (SP 51·16 vs nSP 53·98, p=0.15; Grade IV only: SP 54·07 vs nSP 55.04, p=0.65) or pre-surgery Karnofsky Performance Score (KPS, p=0.21) between the SP and nSP groups. However, after calculating the age-adjusted rate of seizures by using two larger glioblastoma (GB) populations as “standard populations”^18^, we observed younger patients more frequently present with seizures than older patients (more details in **Supplement: Table S1 and Figure S1)**.

### IDH1 Mutation

Since previous literature reported a strong association between seizure presentation in gliomas and IDH1 mutation^10,^ 19, we examined the relationship between this genetic modifier and seizure presentation. IDH1 mutation status was available for 35% of the cohort (78/223; LGG 12/78, 15%), (**Supplement Table S3**). Our cohort (all grades) included 66 IDH1 wild-type (wt) by immunohistochemistry (25 SP and 41 nSP) and 12 IDH1 Mutant (Mut, six SP and six nSP). Among all grades, seizure presentation status was not dependent on IDH1 mutation status (Fisher’s exact test, p=0.53), though the numbers are small. Notably, though not statistically significant, the majority of grade III IDH1Mut patients were SP (83%), while the majority of grade III IDH1wt patients were nSP (75%, Fisher’s exact test, p=0.19). In contrast for Grade IV GB the majority of IDH1wt and IDH1mut were nSP (Fisher’s exact test, p= 0.40).

### Tumor Size and Relative Levels of Invasiveness

Tumor size was significantly smaller in the SP group compared to the nSP group in both T1Gd and T2/FLAIR in the cohort as a whole and when separated by sex (**Figure 1 and Supplement Figure S2**). SP patients had significantly smaller T1Gd radius compared to nSP (All grades: SP 11.30mm ± 6.12, nSP 18.66mm ± 5.88, p<0.0001; Grade IV: SP 12.25mm ± 6.06, nSP 19.32mm ± 5.33, p<0.0001), Similarly, the T2/FLAIR radius was smaller in the SP group compared to the nSP group (All grades: SP 21.54mm ± 7.22, nSP 27.39mm ± 6.94), p<0.0001; Grade IV: SP 21.27mm± 7.24, nSP 27.98mm ± 6.67, p<0.0001). The differences in tumor sizes between the two groups seem to be driven by grade IV gliomas, since grade III differences by seizure presentation group were not significantly different for any of the sizes. There was no statistical difference in tumor invasiveness (log(p/D) between SP and NSP (All grades: SP −0.177± 0.58), nSP −0.059 ± 0.765, p=0.21; Grade IV: SP −0.126 ± 0.545, nSP −0.100 ± 0.701, p=0.79). Importantly, among grade I-IV females, tumors in the SP group (−0.26mm^−2^ ± 0.55) were more diffuse compared to nSP (0.13mm^−2^ ± 0.87).

**Figure 1.**
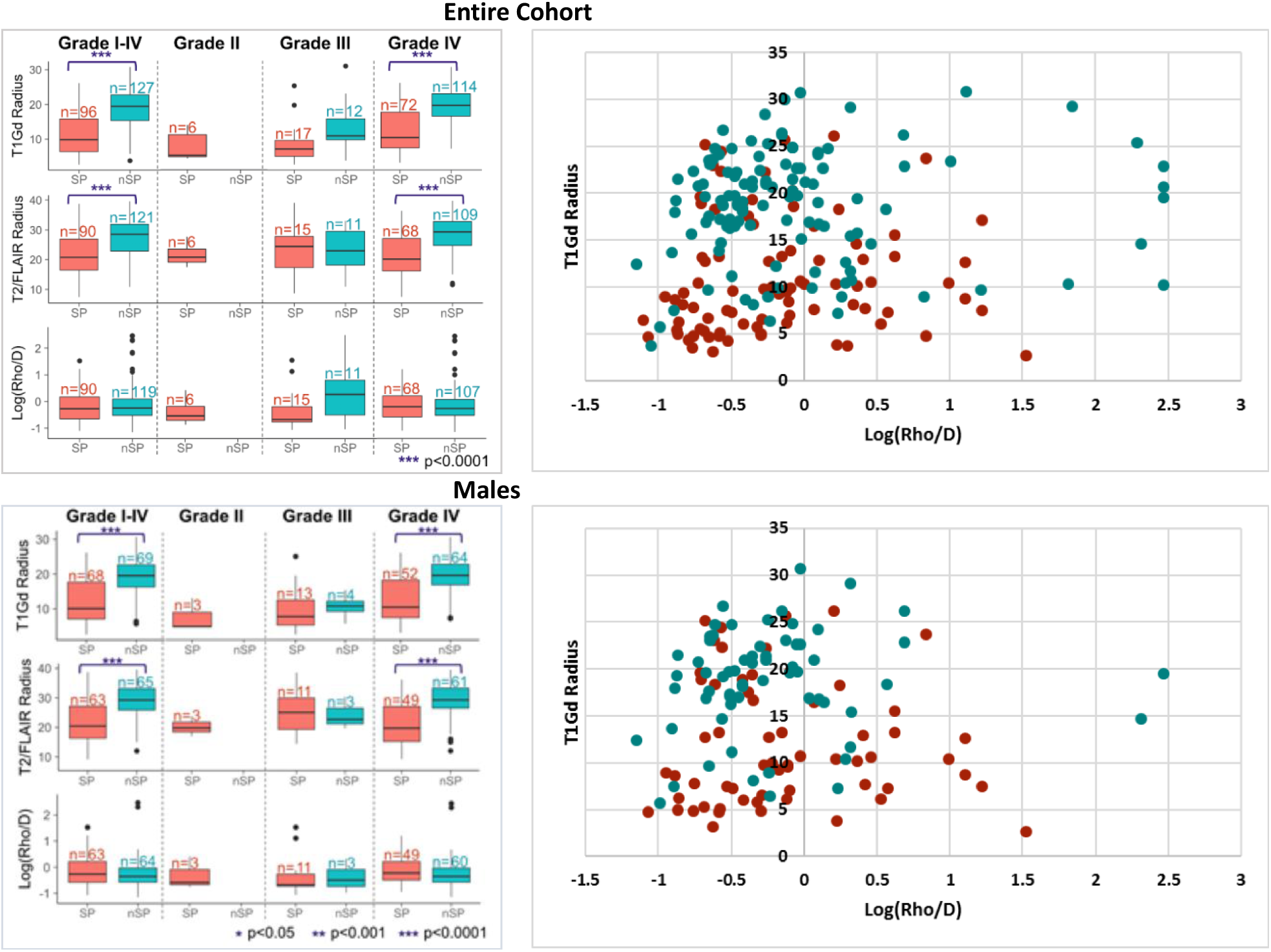
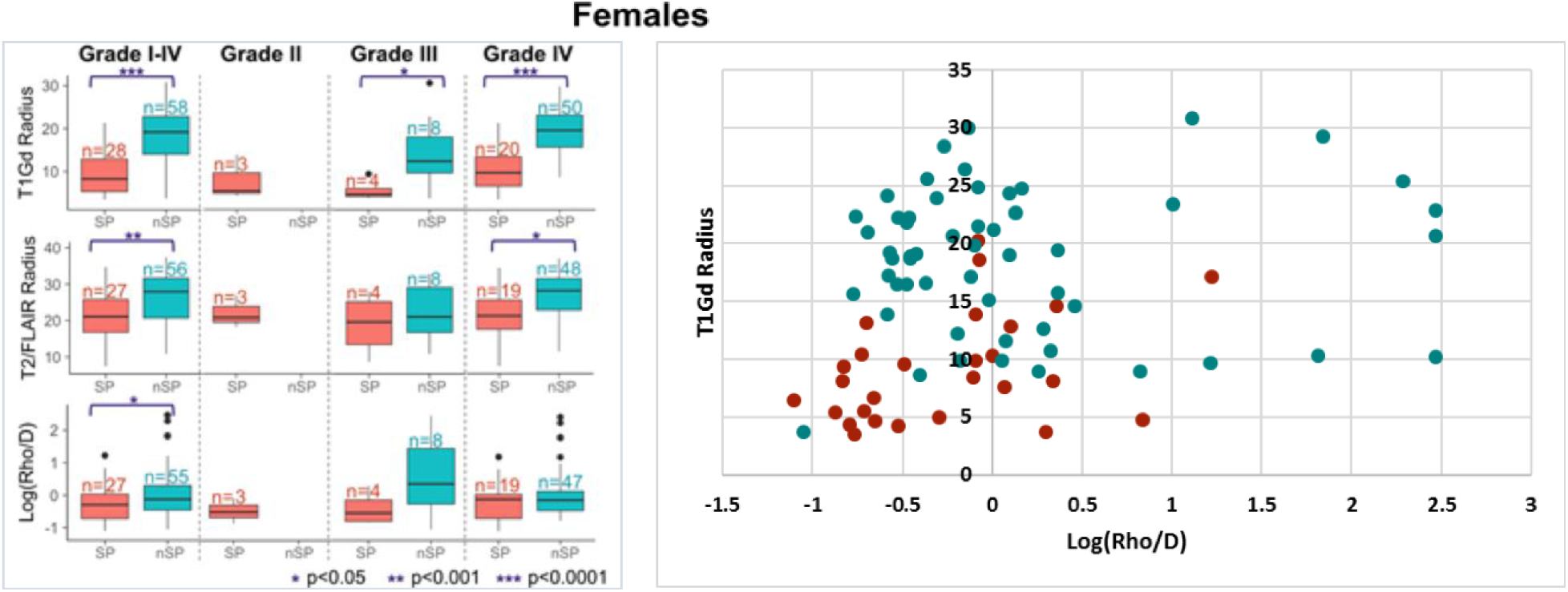
Tumor Size and Invasiveness by Grade and Sex. Comparison of tumor size (left panel as seen on T1Gd above and T2/FLAIR MRI-**Supplement Figure 2**) and relative level of tumor invasiveness by SP (red) and nSP (green) and patient sex. Overall pre-surgery tumor size (T1Gd and T2/FLAIR) are smallest in SP patients, primarily driven by Grade IV GB patients. Relative rates of tumor invasiveness did not differ significantly by seizure presentation status. However, among grade I-IV patients, female SP had significantly more diffuse tumors compared to female nSP. Representation of T1Gd tumor size and degree of tumor invasiveness in a scatterplot (right panel) further illustrating the diffuse tumor associated with SP in females, particularly in smaller tumors (**Supplement Figure S2** tumor invasiveness by FLAIR/T2 and T1Gd and FLAIR/T distribution by sex).

### Tumor location

Seizure presentation status with newly identified gliomas is significantly dependent on tumor location (x^2^=8.04, p=0.05, **Figure 2**). Compared to the percentage of patients presenting with seizure in the entire cohort, tumors in the temporal, frontal, and parietal lobes presented with seizures with quite similar frequency. Compared to all other lobes, patients with tumors in the occipital lobe rarely presented with seizure. While seizure presentation status was not statistically dependent on tumor laterality (x^2^=2.60, p=0.11), we observed that right-sided tumors were predominantly nSP, while left sided tumors were split more evenly between the two groups. The majority of tumors comprised more than one lobe location (multi-lobular: 64%); nSP had a significantly greater portion of multi-lobular tumors (nSP=70%, SP=55%, p=0.004, Fisher’s exact test).

**Figure 2.**
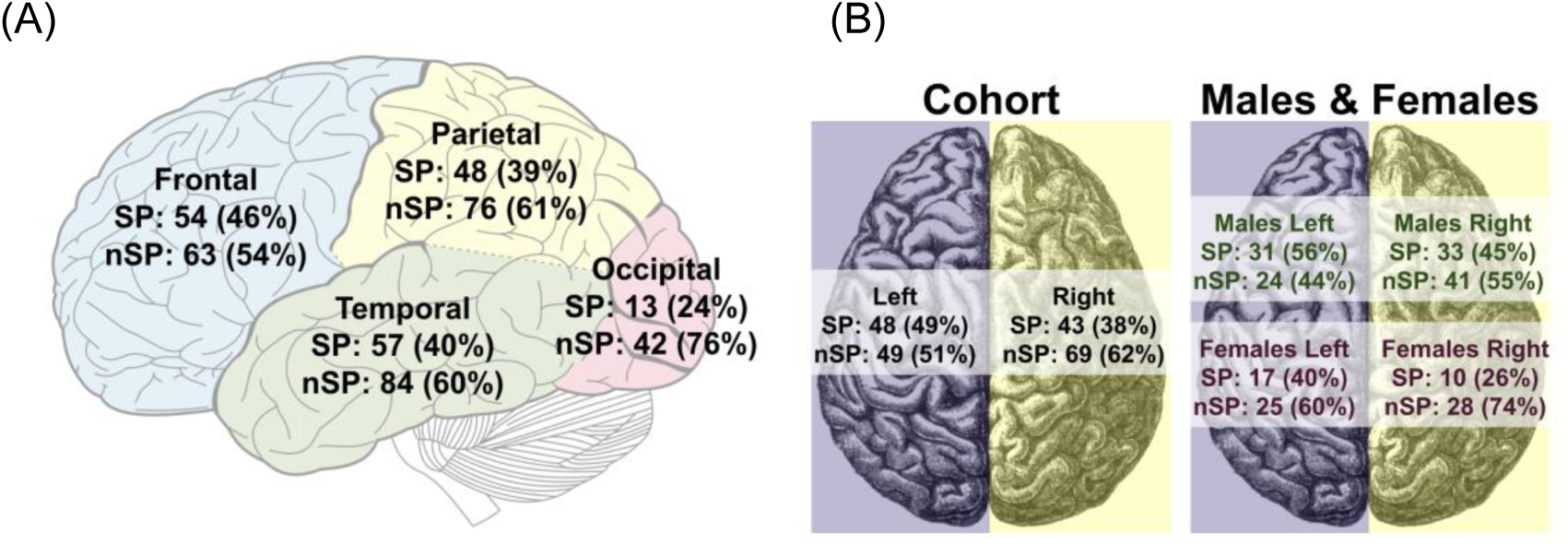
Tumor Location by lobe (A) and hemisphere (B). Distribution of tumors locations were in this order: temporal, frontal, and parietal lobes for both SP and nSP. The predominance of nSP among right-sided tumors seems to be driven by females, with 74% of female right-sided tumors being nSP, while male right-sided were split more evenly between the two groups. Additional sex differences detailed in **Supplement Figure S3**. Images acquired from (A) stock image of brain lobes from this open web source, content added: https://commons.m.wikimedia.org/wiki/File:Lobes_of_the_brain_NL.svg and (B) stock image from https://commons.m.wikimedia.org/wiki/File:Brain_Lateralization.svg under license Creative Commons Attribution-Share Alike 3.0 Unported with color revision and content added over the brain image.

### Survival

Survival statistics are presented as median values from K-M analysis. For the entire cohort, SP patients had longer OS (712 days, IQR 1542) compared to nSP patients (559 days, IQR 598), albeit this difference was not statistically significant (log-rank, p=0.12, Figure 3). Amongst females, the survival difference is larger between SP and nSP, but not statistically significant (**Supplement Figure S3**). However, amongst grade III patients, SP patients had significantly longer OS than nSP (SP OS: 1925 days, IQR 1296; nSP 551 days, IQR 280.5; p=0.004). This difference was not seen amongst grade IV patients (SP 552 days, IQR 947; nSP 559 days, IQR 604.25; p=0.77).

**Figure 3.**
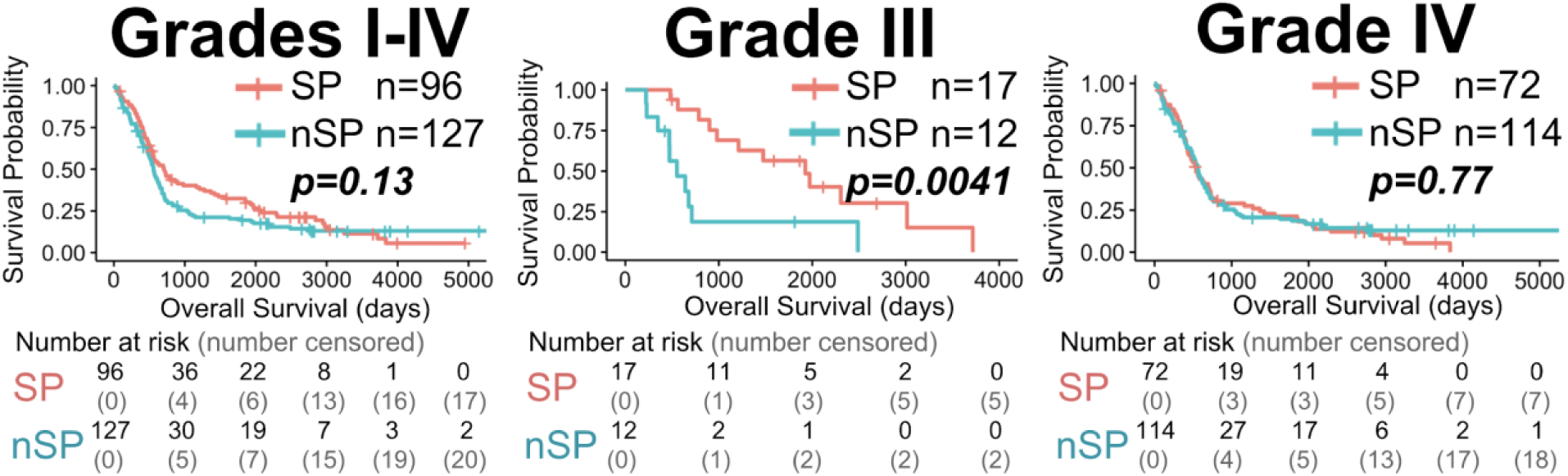
Overall Survival by Tumor Grade and Clinical Seizure Presentation. All SP patients lived longer than nSP but this finding is only statistically significant for Grade III. See **Supplement Figure S4** for OS and tumor grade by patient sex.

### Machine Learning Predictors

**Figure 4** compares the importance of features to each model (A) and presents receiver operating characteristic (ROC) curves of the models in cross-validation (B). The female model achieved the highest AUC (0.853) followed by the mixed cohort with an AUC=0.726. In comparison, the performance of the male-only model was considerably worse (AUC=0.651). **Table 2** shows detailed performance of the three classification models in cross-validation and on previously unseen test sets. Model performance on the test set did not seem to deviate from cross-validation performance for any cohort. **Supplement Table S4** shows the correlation between predicted probability of SP and the continuous numeric variables in the validation data. The Pearson test of correlation showed significant negative correlations (decreased chance of seizure with increase in feature value) between probability of seizure (p(SP)) and T1Gd and T2/Flair in the male and mixed models (p < 0.05). The mixed cohort model had an additional significant negative correlation between+ log(PIHNA D) and p(SP) (p<0.05). In the mixed and male models, smaller tumors seemed to be associated (p<0.05) with higher p(SP). However, the female model did not show any significant correlations with p(SP). Interestingly, males showed a significantly higher predicted probability of seizure for higher tumor invasiveness measures (PI ***ρ***/D). Age had a negative but insignificant correlation with predicted probability of seizure (p<0.1) in the male and female models.

**Figure 4:**
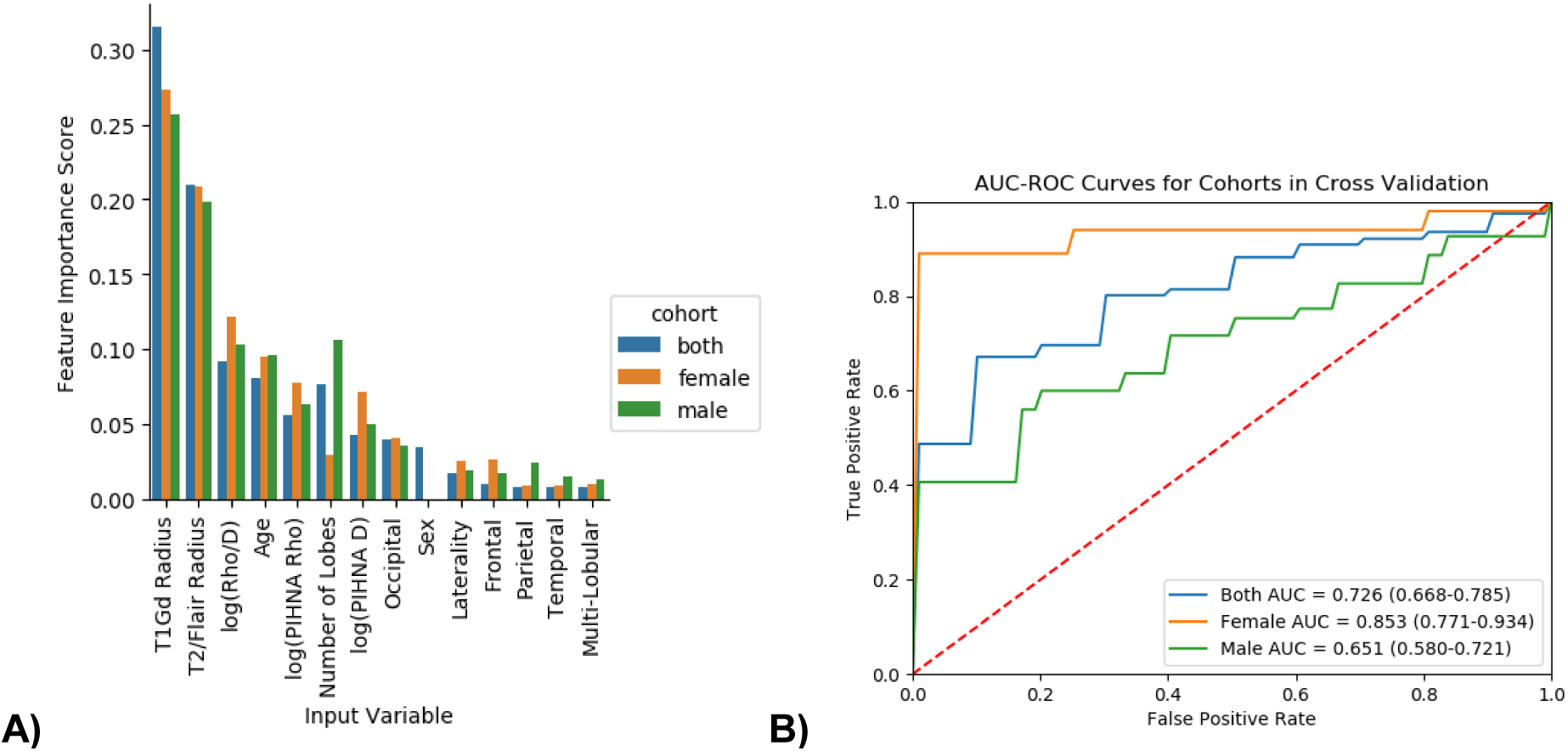
Comparison of Model Performance Among Three Cohorts Grouped by Patient Sex. **A**) The feature importance scores for random forest classifiers trained on each of the cohorts (male, female and combined male, female). The plots demonstrate the statistical prioritization of the variables used in model training. As shown, T1Gd and T2-Flair size had the two highest scores for all three cohorts. For the male-cohort, number of lobes with tumor had the third highest feature importance score, while for the mixed sex (males and females) and female-only cohorts, p/D was the third highest feature importance score. Numeric values for feature importance in the models are presented in **Supplementary Table S5. B**) The ROC curves for the three models. Area under the ROC curve (AUC) as well as confidence intervals are presented in legends. The female-only cohort had the largest AUC of 0.853.

**Table 2.**
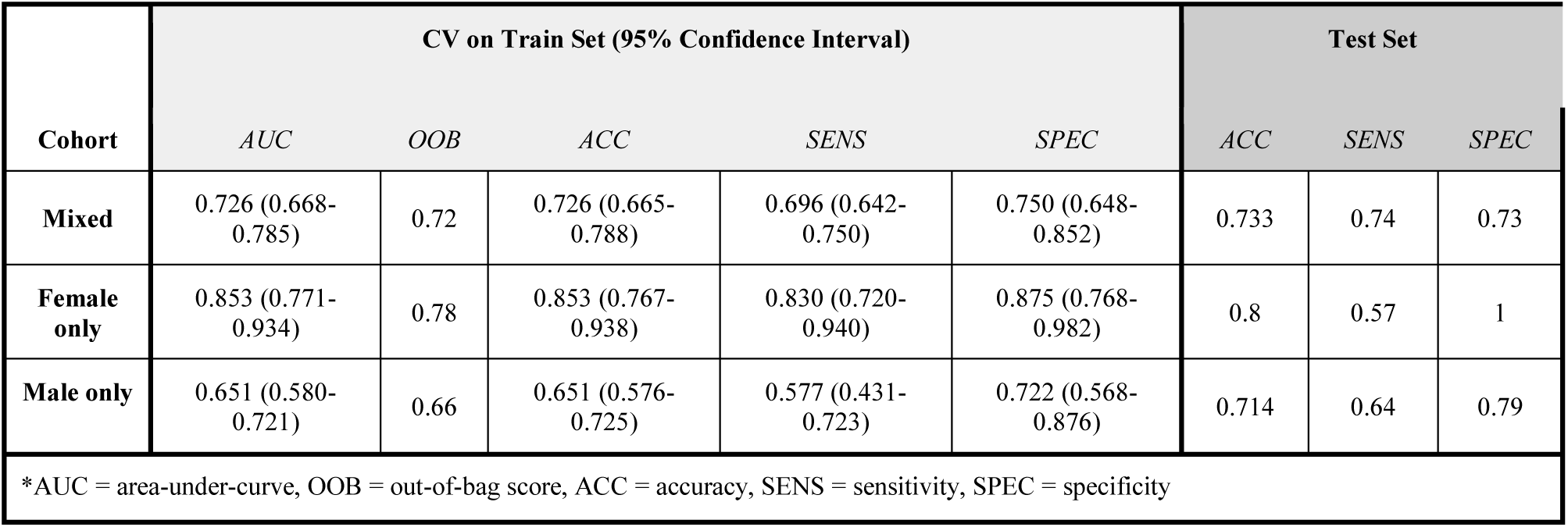
Performance of machine learning in prediction of seizure at presentation for the entire cohort and separated by sex (male, female).

| Cohort | CV on Train Set (95% Confidence Interval) |  |  |  |  | Test Set |  |  |
| --- | --- | --- | --- | --- | --- | --- | --- | --- |
|  | AUC | OOB | ACC | SENS | SPEC | ACC | SENS | SPEC |
| Mixed | 0.726 (0.668-0.785) | 0.72 | 0.726 (0.665-0.788) | 0.696 (0.642-0.750) | 0.750 (0.648-0.852) | 0.733 | 0.74 | 0.73 |
| Female only | 0.853 (0.771-0.934) | 0.78 | 0.853 (0.767-0.938) | 0.830 (0.720-0.940) | 0.875 (0.768-0.982) | 0.8 | 0.57 | 1 |
| Male only | 0.651 (0.580-0.721) | 0.66 | 0.651 (0.576-0.725) | 0.577 (0.431-0.723) | 0.722 (0.568-0.876) | 0.714 | 0.64 | 0.79 |

\*AUC = area-under-curve, OOB = out-of-bag score, ACC = accuracy, SENS = sensitivity, SPEC = specificity

## Discussion

### Sex

Our analysis found that tumor growth kinetics have a different relationship with seizure presentation depending on patient sex. Among males, the rate of invasion was a negative predictor of SP, while the ratio of proliferation to invasiveness was a positive predictor of SP. In females, tumor growth kinetics were not significant predictors of SP. Additionally, the majority of SP were male (70.8%) and a higher proportion of SP males (49.6%) compared to SP females (32.6%). Similarly, Lapointe reported significantly greater number of males in the SP group (66%) and overall larger proportion of SP males (44.6%) than SP females (26.7%)^20^, while others have found no difference in sex distribution^21,22^. These findings are in concert with Christensen’s findings in general epilepsy with males having a slightly higher incidence than females^6^.

### Age

Unlike others^2,20,21,23,24^ we did not find SP patients to be significantly younger than nSP when considering all grades of glioma. However, we did find that SP was more common among younger patients using an age-adjusted rate of SP in grade IV patients.

### Tumor Pathology

The frequency of seizure as an initial symptom of glioma varies across studies and tumor grade. Overall 43% in our cohort were SP well within the range (32% to 86%) reported in the systematic review by Su of all glioma grades^25^. As in our cohort (all six Grade II were SP) others report a greater prevalence of SP in LGG^20,21,23,26^. These LGG tumors are slowing growing which may create areas of cortical denervation hypersensitivity that give rise to seizures. This phenomena was documented in the 1930’s to explain the cause of focal epilepsy^27^. In contrast as fast growing tumors, HGG may affect neurological changes likely through necrosis and hemorrhage^22^. In our cohort of Grade III gliomas, SP was more common in IDH1mut tumors. In contrast, Phan’s systematic review found near equal proportions of IDH1mut and IDH1wt in grade III SP^19^.

### Tumor Location Lobes and Laterality

Tumor location matters in SP, but the methods used to ascertain tumor location strongly vary across groups making comparisons and conclusions difficult. Many^22,25,28,29^ studies reported highest incidences for frontal lobe tumors are associated with SP in both univariate and multivariate analyses. While Lee found temporal lobe tumors in LGG were twice as likely for SP but not true in HGG^24^. This corresponds to the observation that focal, non-tumor seizures most commonly involve the temporal lobe. Our findings are similar to Lynam’s group^26^ with a preponderance of tumors for SP patients located in (in order of occurrence) parietal, temporal, and frontal lobes. Other groups have found no difference in SP status by tumor location^21,23^.

While there are hemispheric links based on the dominant side to cognitive disorders in glioma, seizures in glioma are not similarly associated^23^. As with others^21,23^ we found no association between laterality and seizure presentation. Interestingly, we observed that a greater proportion of right-sided tumors were nSP (62%) compared to SP (38%) while left-sided tumors had equal proportions (SP, 49%, nSP 51%). Similarly, Yang found tumors in the left hemisphere statistically associated with SP^28^.

### Tumor Size

As with prior reports^22,30^, we found smaller contrast-enhancing tumors were significantly associated with SP while accounting for tumor grade. Lee found smaller tumor size associated with HGG but not LGG^24^. Pallud found no relationship with tumor size and seizure presentation in their analysis of LGG^2^. This smaller tumor size in SP alone may help explain the trend toward longer OS in SP.

### Overall Survival

SP was significantly associated with increased OS independent of age, sex, tumor location, and size in the systematic review of Phan 2018^19^. However, we and others found that OS was not dependent on clinical seizure presentation^10,29,30^ despite smaller tumor size in the SP group. We did find that Grade III SP patients had significantly greater OS.

### Machine Learning Predictive Models

To the best of our knowledge, we are the first to report the use of machine learning for identification of clinical factors that could predict seizure at clinical presentation of glioma. Our model shows that at least for the combined male/female cohort and male cohort, tumor size and invasiveness were significant negative predictors for SP; patients in these cohorts with larger tumors and less tumor diffusivity (***ρ***/D) were at lower risk for SP.

### Practice Parameters

AEDs continue to be prescribed in seizure-naïve instances associated with glioma presentation notwithstanding the published practice parameters^1^ that clearly state AEDs are not effective in prevention of first seizure in gliomas providing justification against prophylactic treatments. This fact is reported time and again, most recently in Sirven (meta-analysis) and Lapointe reported, a lack of efficacy for seizure prevention with prophylactic AEDs^12,20^.

Furthermore, AEDs, including new generation drugs, may be associated with serious AEs further supporting the argument again prophylactic use in gliomas. In fact, AEDs complicate oncological treatments with decreased efficacy and potential to compound bone marrow suppression and liver dysfunction. Further, AEDs can decrease cognitive function, an already suppressed function in glioma further negatively impacting QOL.

### Limitations and future opportunities

Reliance on retrospectively available clinical data allowed us to focus on differences in clinical characteristics between SP and nSP patients. Further, we relied upon clinician-derived medical history and not EEGs in determining seizure presentation status. Additional prospective studies are needed in this subject to help guide and clarify AED utilization for clinicians. By identifying some of the clinical and kinetic characteristics associated with seizure presentation, our study is an effort to set up a clearer foundation for these studies.

## Acknowledgements

Cassandra Rickertsen, Gustavo De Leon, Lisa Paulson, Spencer Bayless and all of the many undergraduate students that have worked on the Image Analysis Team in the Dr. Swanson’s Mathematical Neuro-Oncology Laboratory, Precision Neurotherapeutics Innovation Program lab.

