## Supplement for "Sex Differences in Predictors of Seizure in Contrast-Enhancing Gliomas at Clinical Presentation: A Network Approach" (from another version of this preprint: version 1, 2019-07-22)

**Age-adjusted rate of seizure presentation:**

Given that GB incidence varies with age, we compared the number of SP patients in each age group to estimates of GB incidence for that age group in two "standard populations." These comparisons allow us to obtain a more accurate comparison of seizure presentation frequency across age groups. The "age-adjusted SP rate" was calculated as (number of SP in a given age group from our main seizure cohort)/(number of GB patients in that age group in the "standard population"). To ensure validity of results, two "standard populations" were used: 1) the CBTRUS 2011-2015 GB population, and 2) a previously published cohort of 494 first-diagnosis GB patients (Whitmire, 2018) sourced from the same repository as the original seizure cohort. Since we do not have the total SP count from either "standard population", the calculated SP rates are only intended for comparisons across age group within a single "standard population" comparison, not for prediction of seizure frequency. When looking across age groups, both "standard populations" showed a similar pattern of decreasing rate of SP as age increases, suggesting that younger patients more frequently present with seizures compared to older patients.

| **Age Group** | **Number of grade IV SP** | **CBTRUS count* (n=8665)** | **Age-adjusted SP rate (CBTRUS)**** | **494 patient repository cohort** | **Age-adjusted SP rate (repository)***** |
|---|---|---|---|---|---|
| 20-34 | 8 | 255.26 | 3.134 | 28 | 2.857 |
| 35-44 | 6 | 558.24 | 1.075 | 33 | 1.818 |
| 45-54 | 22 | 1314.6 | 1.674 | 104 | 2.115 |
| 55-64 | 20 | 1928.9 | 1.037 | 159 | 1.258 |
| 65-74 | 10 | 2355.8 | 0.4245 | 107 | 0.9346 |
| 75-84 | 6 | 1863.2 | 0.3220 | 45 | 1.333 |
| 85+ | 0 | 386.31 | 0.0000 | 6 | 0.0000 |

**Table S1. Table of data used to calculate age-adjusted SP rate.** Patients under 20 years old in the "standard populations" were excluded from this analysis; all grade IV SP patients were older than 20.

*The number of GB patients in each age group was calculated from the "age-adjusted incidence rate" of GB and 2000 US standard population counts provided in the CBTRUS 2011-2015 report.

**CBTRUS age-adjusted SP rates are represented as SP cases per 100 GB patients (raw rate x 100)

***Repository age-adjusted SP rates are represented as SP cases per 10 GB patients (raw rate x 10)

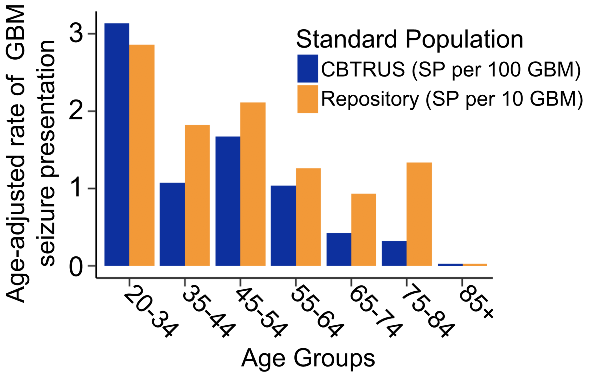


**Supplement Figure S1**. **Age-adjusted rates** of SP among grade IV patients show that seizure presentation is more common among younger patients. Rates are calculated by comparing the number of SP patients from a particular age group in our population to the number of GB cases in that age group in two "standard populations" (CBTRUS 2011-2015 and a GB population from our repository). Rates are intended for comparisons across age group within a single "standard population" comparison, not for prediction of seizure frequency. Rates calculated using both "standard populations" show similar trends of decreasing seizure frequency with increasing age.

### Statistical sex differences analysis:

**Males**

| | Seizure at Presentation (SP) | | | No Seizure at Presentation (nSP) | | | SP vs nSP Statistical Comparison |
|---|---|---|---|---|---|---|---|
| | Mean ± SD | Range | n=___ | Mean ± SD | Range | n=___ | |
| Age at Diagnosis | 52.25 ± 14.07 | 21-80 | 68 | 53.83 ± 13.91 | 19-89 | 69 | p=0.51 (t-test) |
| Pre-surgical KPS | 86.33 ± 12.17 | 40-100 | 30 | 83.97 ± 9.472 | 50-95 | 39 | p=0.38 (t-test) |
| Overall Survival | 1072 ± 1000 | 13-4948 | 68 | 967.2 ± 1137 | 44-5314 | 69 | p=0.40 (log-rank) |
| Tumor Grade | | Grade I: n=0<br>Grade II: n=3<br>Grade III: n=13<br>Grade IV: n=52<br>High-grade: n=0 | | | Grade I: n=1<br>Grade II: n=0<br>Grade III: n=4<br>Grade IV: n=64<br>High-grade: n=0 | | p=0.015<br>$x^2$=5.941<br>(Grade III-IV, chi-square) |

**Females**

| | Seizure at Presentation (SP) | | | No Seizure at Presentation (nSP) | | | SP vs nSP Statistical Comparison |
|---|---|---|---|---|---|---|---|
| | Mean ± SD | Range | n=___ | Mean ± SD | Range | n=___ | |
| Age at Diagnosis | 48.50 ±15.51 | 22-78 | 28 | 54.16 ± 14.96 | 19-75 | 58 | p=0.12 (t-test) |
| Pre-surgical KPS | 86.67 ± 5.000 | 80-90 | 9 | 83.18 ± 11.29 | 60-100 | 22 | p=0.24 (t-test) |
| Overall Survival | 1448 ± 1329 | 92-3996 | 28 | 842.4 ± 950.6 | 13-3897 | 58 | p=0.14 (log-rank) |
| Tumor Grade | | Grade I: n=0<br>Grade II: n=3<br>Grade III: n=4<br>Grade IV: n=20<br>High-grade: n=1 | | | Grade I: n=0<br>Grade II: n=0<br>Grade III: n=8<br>Grade IV: n=50<br>High-grade: n=0 | | p=0.74<br>$x^2$=0.1122<br>(Grade III-IV, chi-square) |

**Supplement Table S2**: Table 1 from main text (Subject demographics and tumor characteristics) separated by sex (male, female). The dependence between seizure presentation status and tumor grade was only found in males and was not statistically significant in females.

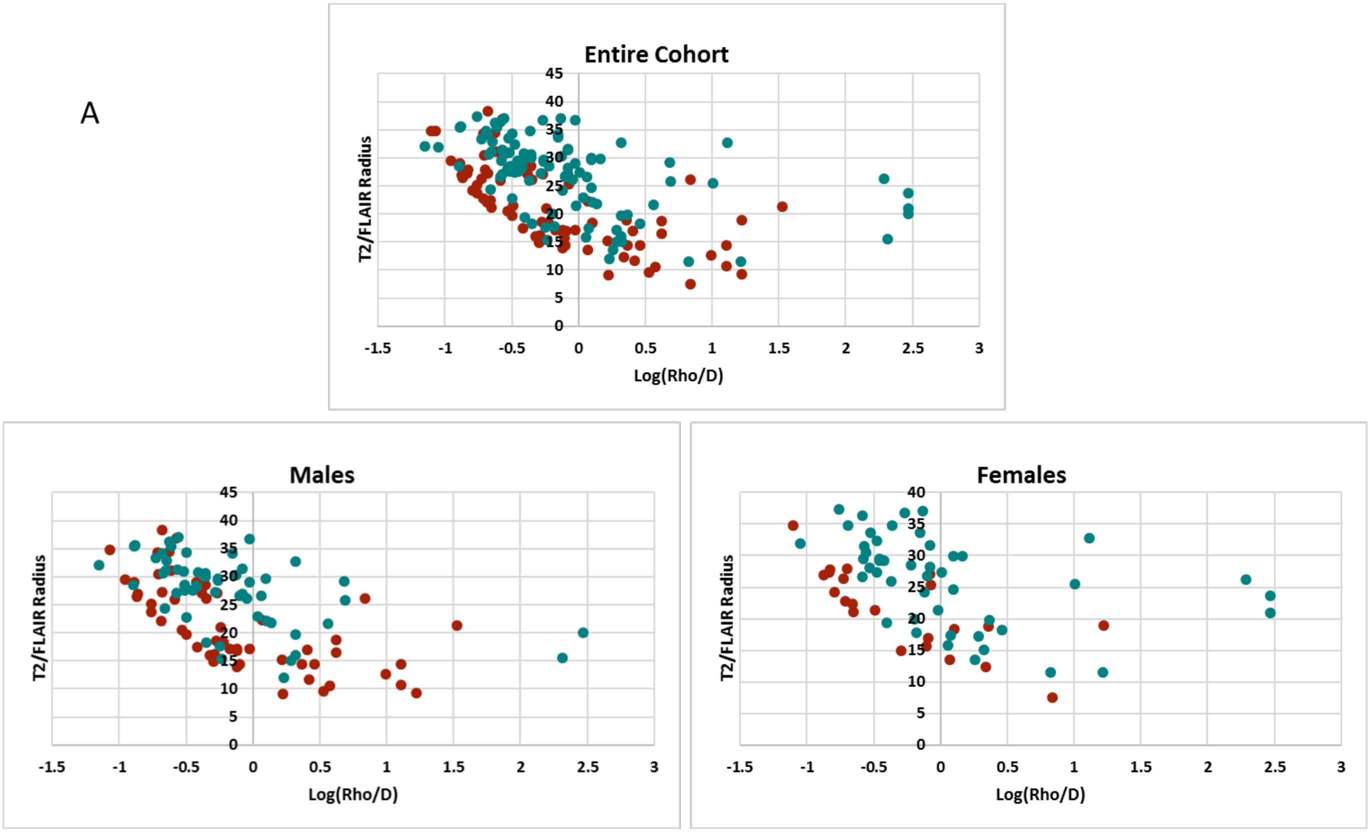

B

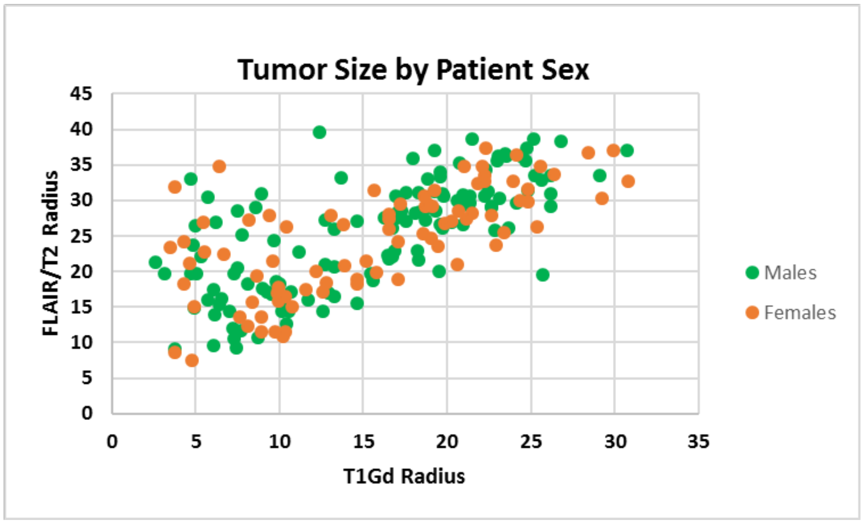


**Supplement Figure S2**. Figure 1 from Main Text (Tumor Size and Invasiveness by Grade and Sex) with FLAIR/T2 separated by sex (male, female) (A), illustrates the diffuse tumor associated with SP in females, particularly in smaller tumors. (B). T1Gd and T2 similarly distributed in males and females.

**Entire Cohort**

| | IDH1 Wild-type | IDH1 Mutant | Fisher's exact test |
|---|---|---|---|
| Grade I-IV | 25 SP (38%)<br>41 nSP (62%) | 6 SP (50%)<br>6 nSP (50%) | p=0.53 |
| Grade II | 0 SP<br>0 nSP | 0 SP<br>0 nSP | N/A |
| Grade III | 1 SP (25%)<br>3 nSP (75%) | 5 SP (83%)<br>1 nSP (17%) | p=0.19 |
| Grade IV | 24 SP (39%)<br>38 nSP (61%) | 1 SP (17%)<br>5 nSP (83%) | p=0.40 |

**Males**

| | IDH1 Wild-type | IDH1 Mutant | Fisher's exact test |
|---|---|---|---|
| Grade I-IV | 19 SP (43%)<br>25 nSP (57%) | 6 SP (60%)<br>4 nSP (40%) | p=0.49 |
| Grade II | 0 SP<br>0 nSP | 0 SP<br>0 nSP | N/A |
| Grade III | 1 SP (50%)<br>1 nSP (50%) | 5 SP (83%)<br>1 nSP (17%) | p=0.46 |
| Grade IV | 18 SP (43%)<br>24 nSP (57%) | 1 SP (25%)<br>3 nSP (75%) | p=0.63 |

**Females**

| | IDH1 Wild-type | IDH1 Mutant | Fisher's exact test |
|---|---|---|---|
| Grade I-IV | 6 SP (27%)<br>16 nSP (73%) | 0 SP (0%)<br>2 nSP (100%) | p=1.0 |
| Grade II | 0 SP<br>0 nSP | 0 SP<br>0 nSP | N/A |
| Grade III | 0 SP (0%)<br>2 nSP (100%) | 0 SP<br>0 nSP | p=1.0 |
| Grade IV | 6 SP (30%)<br>14 nSP (70%) | 0 SP (0%)<br>2 nSP (100%) | p=1.0 |

**Supplement Table S3**: **IDH1 mutation status by seizure presentation**. This is for entire cohort and separated into male and female patients. In likeness to the entire cohort described in the main text, neither males nor females show any significant differences in seizure presentation by IDH1 status.

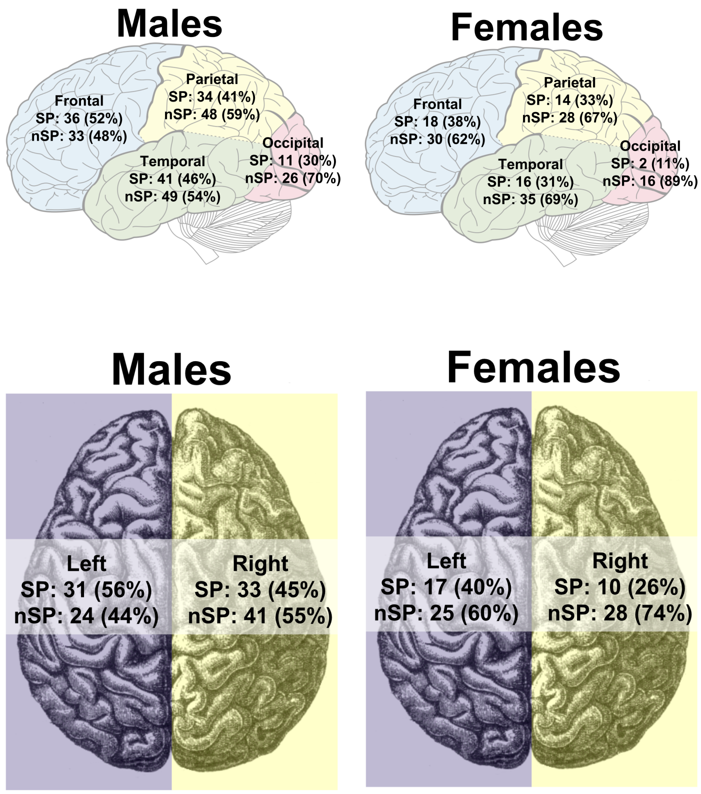


**Supplement Figure S3**: From Main Text Figure 2 (**Tumor Location by Lobe and Hemisphere) with additional sex differences detailed**. Seizure presentation status is not dependent on laterality for males ($x^2$=1.75, p=0.19) or females ($x^2$=1.79, p=0.18). While seizure status is dependent on location for the combined cohort, it is not dependent on location for males ($x^2$=5.23, p=0.16) or females ($x^2$=4.34, p=0.23) separately.

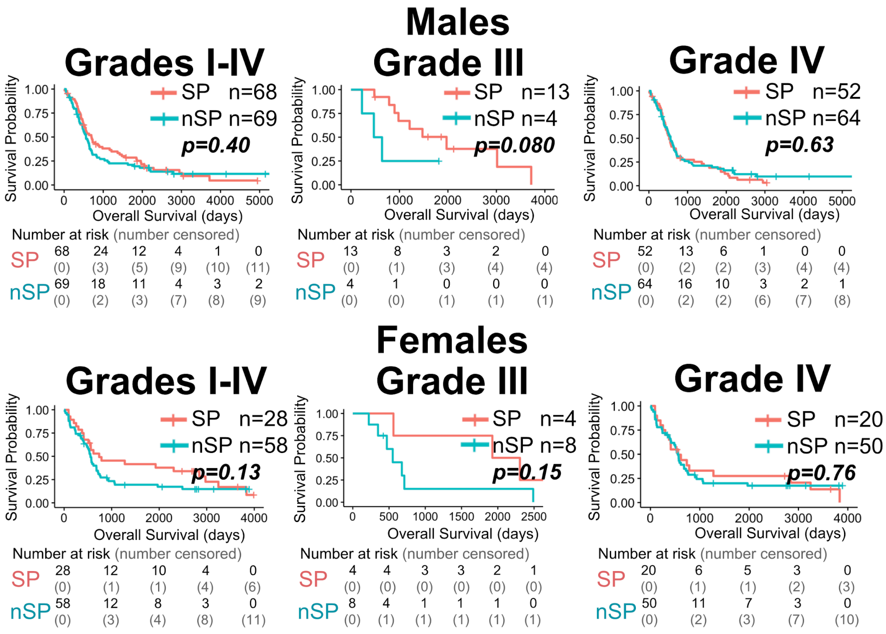


**Supplement Figure S4:** Main Text Figure 3: **Overall Survival by Tumor Grade separated by Sex (male, female)**. Results for males and females look similar to the combined cohort, with the exception that there is more of a difference in survival between SP and nSP for females, although it is not statistically significant.

### Machine learning predictors:

| Attribute | Entire Cohort | Female | Male |
|---|---|---|---|
| T1Gd | -0.41* | -0.06 | -0.40* |
| T2/Flair | -0.38* | 0.045 | -0.45* |
| log(PI ρ/D) | -0.026 | 0.025 | 0.22* |
| log(PIHNA ρ) | -0.059 | 0.01 | 0.035 |
| log(PIHNA D) | -0.19* | 0.01 | -0.18** |
| Age | -0.064 | -0.19** | -0.17** |

**Supplement Table S4 - Correlations between predicted probability of seizure and the continuous variables.** *p-value < 0.05 for Pearson test of correlation; **p-value < 0.1 for Pearson test of correlation

| FEATURE | Entire Cohort | Female | Male |
|---|---|---|---|
| T1Gd Radius | (1) 0.32 | (1) 0.27 | (1) 0.256 |
| T2 Flair Radius | (2) 0.21 | (2) 0.21 | (2) 0.2 |
| $\rho$/D | (3) 0.092 | (3) 0.12 | (4) 0.10 |
| Age | (4) 0.081 | (4) 0.095 | (5) 0.096 |
| PIHNA p (1/yr) | (5) 0.057 | (5) 0.078 | (6) 0.063 |
| Number of sites | (6) 0.077 | (8) 0.030 | (3) 0.11 |
| PIHNA D ($mm^2$/yr) | (7) 0.043 | (6) 0.072 | (7) 0.050 |
| Occipital | (8) 0.040 | (7) 0.041 | (8) 0.035 |
| Sex | (9) 0.035 | (14) 0 | (14) 0 |
| Laterality | (10) 0.017 | (10) 0.026 | (10) 0.020 |
| Frontal | (11) 0.011 | (9) 0.026 | (11) 0.018 |
| Parietal | (12) 0.0078 | (13) 0.0087 | (9) 0.025 |
| Temporal | (13) 0.0081 | (12) 0.0093 | (12) 0.015 |
| Multi-Lobular | (14) 0.0081 | (11) 0.011 | (13) 0.013 |
| **OOB** | **0.719** | **0.776** | **0.662** |

Table S5- **Importance of Features to Predictive Modeling** in the 3 Cohorts (Entire Cohort, Female, and Male)

### SUPPLEMENT REFERENCES

1. Whitmire, P, Rickertsen CR, Hawkins-Daarud, A, Carrasco E, Lorence, J, Swanson, KR et al Sex-specific impact of patterns of imageable tumor growth on survival of primary glioblastoma patients. bioRxiv doi: 10.1101/325464
